# Identification of molecular candidates which regulate calcium-dependent CD8^+^ T-cell cytotoxicity

**DOI:** 10.1101/2020.12.22.423945

**Authors:** Sylvia Zöphel, Gertrud Schwär, Maryam Nazarieh, Verena Konetzki, Cora Hoxha, Eckart Meese, Markus Hoth, Volkhard Helms, Mohamed Hamed, Eva C. Schwarz

**Affiliations:** Biophysics, Center for Integrative Physiology and Molecular Medicine, School of Medicine, Saarland University, Building 48, 66421 Homburg, Germany; Center for Bioinformatics, Saarland University, 66041 Saarbrücken; Human Genetics, School of Medicine, Saarland University, Building 60, 66421 Homburg, Germany; Institute for Biostatistics and Informatics in Medicine and Ageing Research, Rostock University Medical Centre, 18057 Rostock, Germany

**Keywords:** Calcium, CTL (cytotoxic T lymphocytes), real-time killing assay, cytotoxic efficiency, differential expression analyses, transcriptome data analyses

## Abstract

Cytotoxic CD8^+^ T lymphocytes (CTL) eliminate infected cells or transformed tumour cells by releasing perforin-containing cytotoxic granules at the immunological synapse. The secretion of such granules depends on Ca^2+^-influx through store operated Ca^2+^ channels, formed by STIM-activated Orai proteins. Whereas molecular mechanisms of the secretion machinery are well understood, much less is known about the molecular machinery that regulates the efficiency of Ca^2+^-dependent target cell killing. Here, we isolated total RNA from natural killer (NK) cells, non-stimulated CD8^+^ T-cells, and from Staphylococcus aureus enterotoxin A (SEA) stimulated CD8^+^ T-cells (SEA-CTL) and conducted whole genome expression profiling by microarray experiments. Based on differential expression analysis of the transcriptome data and analysis of master regulator genes, we identified 31 candidates which potentially regulate Ca^2+^-homeostasis in CTL. To investigate a putative function of these candidates in CTL cytotoxicity, we transfected either SEA-stimulated CTL (SEA-CTL) or antigen specific CD8^+^ T-cell clones (CTL-MART-1) with siRNAs specific against the identified candidates and analyzed the killing capacity using a real-time killing assay. In addition, we complemented the analysis by studying the effect of inhibitory substances acting on the candidate proteins if available. Finally, to unmask their involvement in Ca^2+^ dependent cytotoxicity, candidates were also analyzed under Ca^2+^-limiting conditions. Overall, this strategy led to the identification of KCNN4, RCAN3, CCR5 and BCL2 as potential candidates to regulate the efficiency of Ca^2+^-dependent target cell killing.

## Introduction

Cytotoxic CD8^+^ T lymphocytes (CTL) and natural killer (NK) cells eliminate infected or transformed tumour cells to control the integrity of the human body. Their main cytotoxic mechanism is the release of perforin/granzyme-containing lytic granules (LG) at the immunological synapse. LG release requires Ca^2+^ entry across the plasma membrane by STIM-activated Orai channels (Maul-Pavicic *et al*., 2011). Beside these key players, Ca^2+^ transporters and other ion channels such as TRPs, CaV, P2X receptors and Piezo channels contribute to the regulation of Ca^2+^-dynamics and Ca^2+^-homeostasis inside cells (Feske *et al*., 2012; Friedmann *et al*., 2019; Trebak & Kinet, 2019). However, their roles for killer cell functioning are not completely understood and controversially discussed (Badou *et al*., 2013; Feske, 2013; Pelletier & Savignac, 2013; Nohara *et al*., 2015; Fenninger & Jefferies, 2019). The function of STIM and Orai proteins is, in principle, well understood and originated from data on SCID patients, whereas our understanding of further proteins often relies on experimental data from human T-cell lines, mice or various other cell lines. We have recently shown that there is an optimal Ca^2+^ concentration for primary human CTL-killing (approximately 25-600 µM of external Ca^2+^ concentration) (Zhou *et al*., 2018) that is much below the free Ca^2+^ concentration in peripheral blood (around 1.3 mM). However, the molecular machinery behind the optimal Ca^2+^ concentration that regulates Ca^2+^ dependent cytotoxicity remains unknown.

In the last 15 years, small interfering RNA (siRNA) based screening strategies have advanced significantly and the therapeutic use of siRNA has been pushed forward as well (Moffat & Sabatini, 2006; Zhou *et al*., 2019; Hattab & Bakhtiar, 2020; Mainini & Eccles, 2020). In primary human cells, RNAi transfection in naïve B-cells results in knockdown efficiencies of about 40–60% (Shih *et al*., 2019). Also, optimized protocols for other primary immune cells (T-cells, monocytes, dendritic cells) were published (Mantei *et al*., 2008; Smith *et al*., 2016; Sioud, 2020). Numerous studies reported successful screens for the function of immune killer cells by siRNA-based approaches. However, the important issue of tumour resistance was often addressed by using either cell lines or transfected target cells (Bellucci *et al*., 2012; Khandelwal *et al*., 2015). Zhou *et al*. successfully discovered negative regulators of T-cell function in the tumour environment based on an in vivo pooled short hairpin RNA (shRNA) screen carried out in OT-I T-cells (Zhou *et al*., 2014).

Detailed analyses of CTL-killing efficiency are of high interest considering the large number of ongoing and upcoming studies on CTL modified for clinical use (i.e., with chimeric antigen receptor T (CAR-T) cells). As the CTL-killing efficiency is Ca^2+^ dependent, it appeared worthwhile to us to knockdown Ca^2+^ related proteins in primary human CTL by siRNAs.

This study aimed at identifying candidates that regulate Ca^2+^ dependent cytotoxicity. For this, genes showing differential expression between NK cells and CTL were intersected with members of known Ca^2+^-signaling pathways and combined with putative key regulators. Subsequently, the effects of the candidates genes were tested after siRNA-silencing them in a real-time killing assay that is independent from the availability of blocking or activating agents.

## Results

Lytic granule (LG)-dependent cytotoxicity of CTL and NK cells depends on Ca^2+^ influx through STIM-activated Orai channels (Maul-Pavicic *et al*., 2011; Zhou *et al*., 2018). Although CTL and NK cells use similar mechanisms to kill their respective targets, the Ca^2+^ dependence of their cytotoxic efficiency is slightly different (Figure 1 and (Zhou *et al*., 2018)). Thus, we were interested in identifying proteins besides STIM and Orai which might have an impact on the regulation of Ca^2+^-dependent killing mechanisms. In the following, these will be termed “candidates”. To this end, we performed expression analysis based on microarray technology of the following human cell populations: 1. un-stimulated CD8^+^ T-cells (purified from human PBMC without any additional cell cultivation, n=11), 2. SEA stimulated CD8^+^ T-cells (SEA-CTL, n=11), 3. NK cells (n=11). Expression data from microarray were background corrected and expression data lower than 2^7.2^ (fluorescence value < 147) were considered as too low or not expressed as confirmed for a number of genes by quantitative RT-PCR.

**Figure 1:**
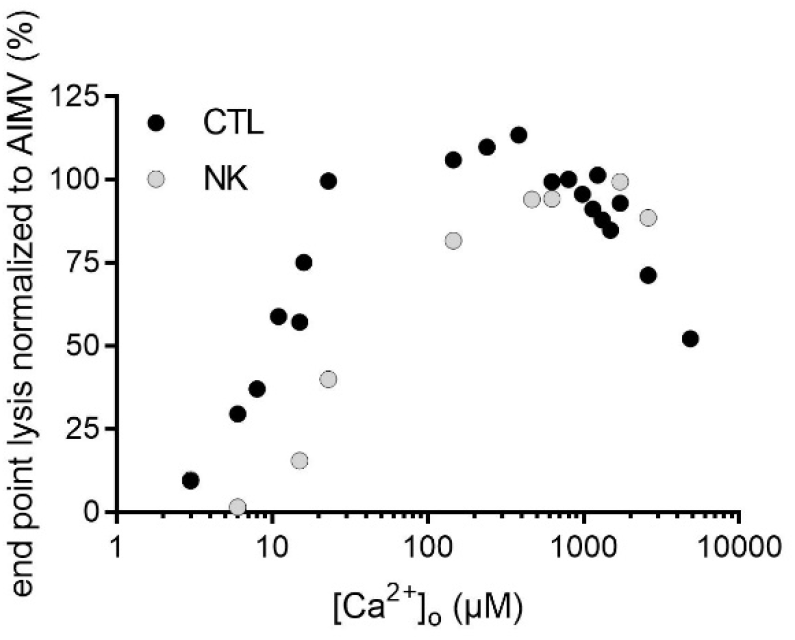
Natural killer cells (NK cells) and cytotoxic T-cells (SEA-CTL) need various external Ca^2+^ concentrations to kill their target cells most efficiently. Data are taken out of Figure 1G and Figure 2H from Zhou *et al*., 2018 and plotted here as a new graph (Zhou *et al*., 2018).

Candidates were selected by two bioinformatic strategies. The first strategy identified 2349 differentially expressed genes (DEG) among NK cells and SEA-CTL from the microarray data. Besides, we identified 512 genes in Ca^2+^-signaling pathways of different databases (for details see Material and Methods). Intersection of these two gene sets gave 86 candidates. Among these 86 candidates, we identified those which were also differentially expressed between un-stimulated CD8^+^ T-cells and SEA-CTL assuming that these genes should play a more prominent role for the cytotoxic functions of stimulated CD8^+^ T-cells. Thereby, 25 genes were eliminated from the 86 candidates. Among the remaining 61 genes, only 29 were reasonably high expressed in SEA-CTL. Thus, this first strategy led to 29 candidates potentially regulating Ca^2+^ dependent CTL cytotoxicity (Figure 2A).

**Figure 2:**
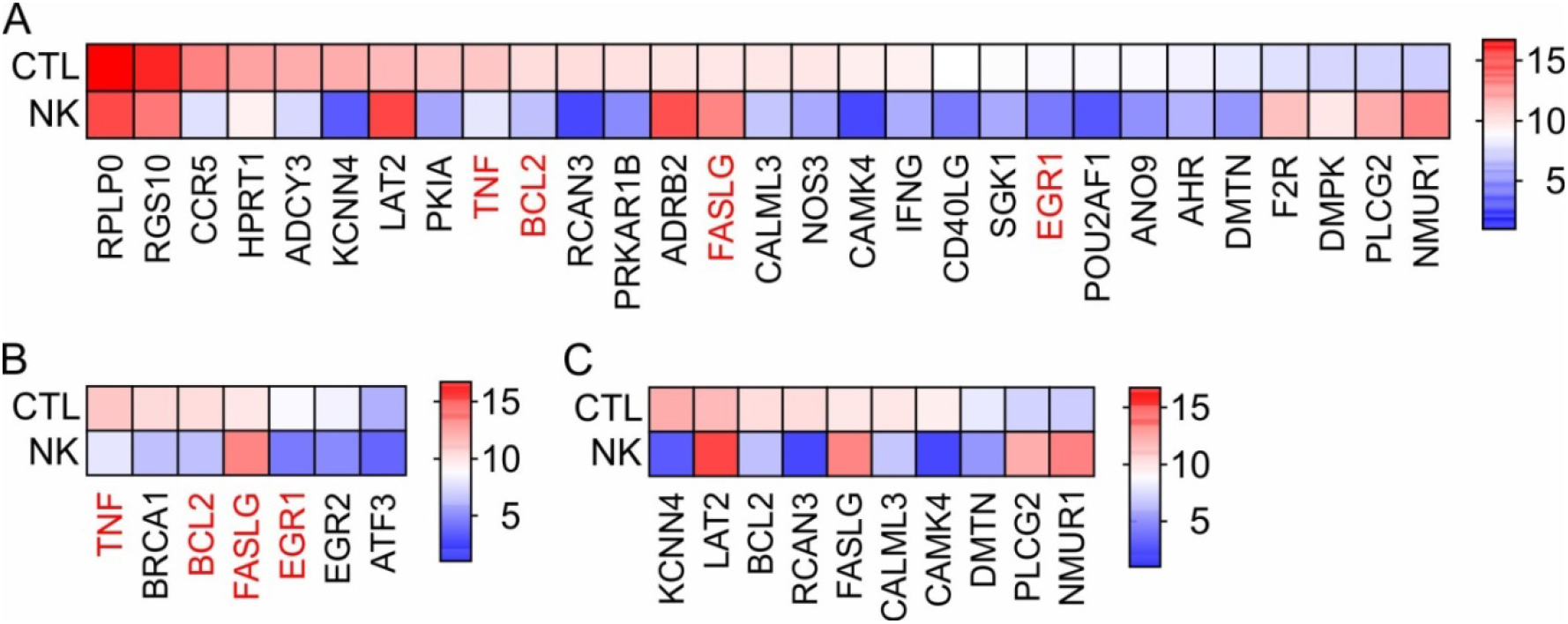
Heatmap representing the expression levels of candidates in SEA-stimulated CD8^+^ T-cells (CTL) and NK cells (NK) identified by bioinformatic analyses. (A) The 29 candidate genes tested in SEA-CTL and CTL-MART-1 identified by strategy 1. (B) The 7 candidate genes identified by strategy 2 and representing Ca^2+^-associated top control genes. Genes labelled in red were identified by both strategies (A,B). C. Candidates randomly selected from all candidates in order to establish our experimental screening strategy in SEA-CTL. The colour scale bars show the corresponding normalized expression values. Blue colour indicates low expression whereas red colour indicates high expression.

In a second strategy, using the prioritization approach TopControl (Nazarieh & Helms, 2019), we prioritized another set of candidates based on a gene-regulatory network of the 2263 DEGs between SEA-CTL and NK cells that was constructed with the TFmiR webserver (Hamed *et al*., 2015). As candidates, we considered hub genes and dominating genes of the regulatory network (see methods). This strategy identified 7 candidates ((Nazarieh & Helms, 2019), Figure 2B). Of these 7 candidates, ATF3 was too low expressed in SEA-CTL, and 4 further candidate genes (BCL2, TNF, FASLG, and EGR1) overlapped with those identified by the first strategy (Figure 2B, labeled in red, Supplementary Figure 1). Combining the candidates from both analyses led to 31 potential candidates.

Before analyzing all 31 candidates, we randomly selected 10 candidates (Figure 2C) to establish our experimental set-up. First, we tested which of the candidates modulate the cytotoxic efficiency in SEA-CTL by down-regulating candidate expression levels via siRNA and a subsequent real-time killing assay (Kummerow *et al*., 2014). As positive control, we transfected an siRNA against perforin which reduced the cytotoxic efficiency as expected (Figure 3A-D, (Zhou *et al*., 2018)). Next, we transfected the SEA-CTL with corresponding siRNAs against the candidates and analyzed these cells 24 and 48 hours after transfection by the real-time killing assay with SEA-loaded Raji cells as targets (Figure 3E). As control conditions, we routinely used SEA-CTL that were each transfected with two different control siRNAs (table 1). All results are always normalized relative to these two controls. The killing efficiency in the perforin siRNA transfected SEA-CTL was reduced by about 42% after 24 hours (perforin mean 26.1% +/−3.4% SEM and ctrl mean 45.0% +/−7.3% SEM) and by about 45% after 48 h (perforin mean 40.2% +/−5.5% SEM and ctrl mean 72.7% +/−9.3% SEM) as shown in representative killing traces (Figure 3A,B) and the endpoint target lysis analyzed in 5-7 different donors (Figure 3C). Furthermore, the maximal killing rates (highest target lysis between two measurement points (%/10 minutes)) confirmed that a stronger effect exists 48 hours after transfection (Figure 3D, 42.5% after 24 hours, 51.7% after 48 hours, reduction compared to control conditions). When we explored the influence of the randomly selected candidates on the killing efficiency, no significant difference to control siRNA transfected SEA-CTL was detected within the sensitivity limit of the killing assay (target lysis at 240 minutes should be either lower than 85% or higher than 115% compared to control transfected siRNA, Figure 3E). Downregulation of several candidates such as KCNN4, BCL2 and DMTN showed a tendency towards a reduced killing efficiency (KCNN4 91.5% relative to ctrl +/−3.9% SEM, BCL2 91.5% relative to ctrl +/−6.5% SEM and DMTN 92.2% relative to ctrl +/−5.7% SEM), but the variability between different experiments was too large to reach statistical significance. Since the observed variability in the killing efficiency of SEA-CTL might unfavourably compromise the interpretation of the results of our initial screen, we implemented another strategy to stimulate CTL. To this end, we used MART-1-specific CTL clones from primary human PBMC (CTL-MART-1, recently established in our lab (Friedmann *et al*., 2020)) and compared the variance in target lysis of CTL-MART-1 to SEA-CTL transfected with control siRNA in the real-time killing assay. Reassuringly, the target lysis at 240 min, analyzed at days 1 and 2 after transfection, was much more stable in CTL-MART-1 than in SEA-CTL (Figure 4A). The cytotoxic efficiency was higher in CTL-MART-1 but most importantly the standard deviation was much lower: Mean and standard deviations of the target lysis after 240 min for MART-1-CTL were 64.0 +/−14.1% (day1) and 68.3 +/−10.9% (day2) compared to 36.3% +/−19.7% (day1) and 53.3% +/−31.2% (day2) for SEA-CTL. The higher variation of SEA-CTL likely results from blood donor variability. In addition to the lower variation of CTL-MART-1, we also found that siRNA down-regulation was more effective in CTL-MART-1 compared to SEA-CTL as tested for 8 candidates (Figure 4B). This was true for all candidates except for BCL-2.

**Figure 3:**
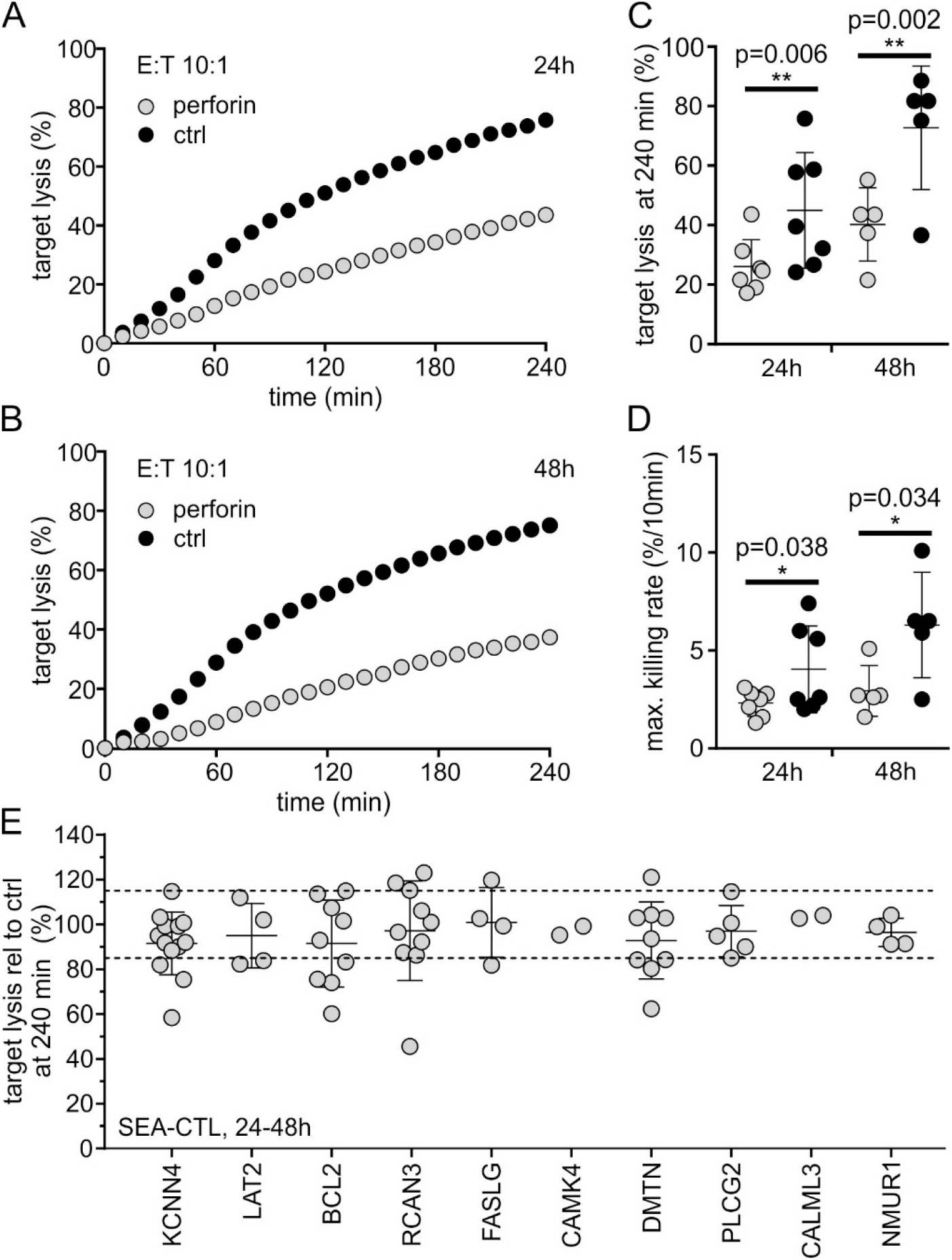
Real-time killing assay of SEA-stimulated CTL (SEA-CTL) as effector cells and SEA-pulsed Raji cells as target cells with an effector to target ratio (E:T) of 10:1 or 20:1. A–D. Real-time killing assay with perforin siRNA transfected SEA-CTL (grey) or the mean of two different control siRNAs (black). A,B. Representative killing traces 24h (A) and 48h (B) after transfection. C,D. Statistical analysis of endpoint lysis after 240 min (C) and maximal killing rates (D), n=7 (24 h after transfection) and n=5 (48 h after transfection). E. Endpoint target lysis of SEA-CTL transfected with siRNAs against 10, randomly chosen candidates (compare Figure 2C) relative to the mean of SEA-CTL transfected with two different control siRNAs (24h and 48h after transfection in AIMV medium). Data are shown as mean +/−SD from 1-10 different donors. KCNN4, n=10; LAT2, n=4; BCL2, n=6; RCAN3, n=5; FASLG, n=4; CAMK4, n=2; DMTN, n=9; PLCG2, n=5; CALML3, n=2; NMUR1, n=4.

**Table 1:** siRNAs used in this study: Perforin siRNA was synthesized by Microsynth sense 5’ OMeA-OMeA (AGAUUGGAUACGCAU)_15_ d(CAG) dOMeA-dOMeT-dOMeT 3’; antisense 3’ OMeT-OMeT-UCUAACCUAUGCGUAGUC-dU 5’.

| gene | gene accession number | Dharmacon, SmartPool order number |
| --- | --- | --- |
| RPLP0 | NM_001002 | L-010864-00 |
| RGS10 | NM_002925 | L-008472-02 |
| CCR5 | NM_000579 | L-004855-00 |
| HPRT1 | NM_000194 | L-008735-00 |
| ADCY3 | NM_004036 | L-006799-00 |
| PKIA | NM_181839 | L-012321-00 |
| TNF | NM_000594 | L-010546-00 |
| PRKAR1B | NM_001164759 | L-030847-02 |
| ADRB2 | NM_000024 | L-005426-01 |
| NOS3 | NM_000603 | L-006490-00 |
| IFNG | NM_000619 | L-019379-00 |
| CD40LG | NM_000074 | L-011006-00 |
| SGK1 | NM_005627 | L-003027-00 |
| POU2AF1 | NM_006235 | L-020133-00 |
| ANO9 | NM_001012302 | L-026726-02 |
| AHR | NM_001621 | L-004990-00 |
| F2R | NM_001992 | L-005094-00 |
| DMPK | NM_004409 | L-004637-00 |
| BRCA1 | NM_007298 | L-003461-00 |
| EGR1 | NM_001964 | L-006526-00 |
| EGR2 | NM_000399 | L-006527-00 |
| LAT2 | NM_014146 | L-013570-02 |
| KCNN4 | NM_002250 | L-004461-00 |
| BCL2 | NM_000657 | L-003307-00 |
| RCAN3 | NM_013441 | L-020467-02 |
| FASLG | NM_000639 | L-011130-00 |
| CALML3 | NM_005185 | L-019122-02 |
| CAMK4 | NM_001744 | L-004944-00 |
| DMTN | NM_001114139 | L-003663-02 |
| PLCG2 | NM_002661 | L-008339-02 |
| NMUR1 | NM_006056 | L-005590-00 |
| non-targ#1 | no accession | D-001210-01 |
| non-targ#2 | no accession | D-001210-02 |

**Figure 4:**
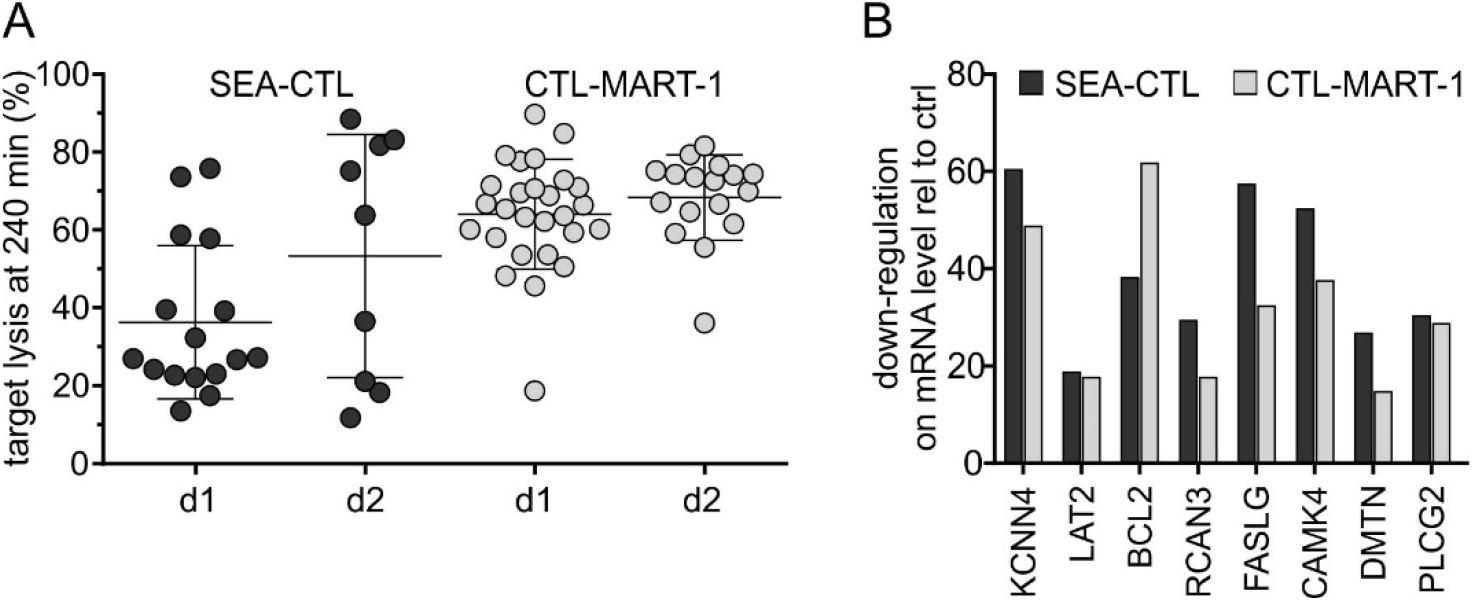
A. Endpoint target lysis of SEA-stimulated CTL (SEA-CTL, black) in comparison to MART-1 specific CTL (CTL-MART-1, grey) one day (d1) or two days (d2) after transfection with control siRNAs. Data points correspond to the mean of two different control siRNAs. Bars show the mean +/−SEM, SEA-CTL d1 n=16, d2 n=9; CTL-MART-1 d1 n=20, d2 n=17. B. Relative mRNA expression of 8 selected candidates 24-30h after transfection of SEA-CTL (black; n=3 for KCNN4; n=4 for BCL2, all other candidates n=1) or CTL-MART-1 (grey; n=3 for KCNN4 and BCL2; n=2 for LAT2; all other candidates n=1) with the corresponding siRNA. The expression level of mRNA in control siRNA transfected SEA-CTL was set to 100%.

Based on the lower variability and the overall better efficiency of down-regulation by siRNA, we switched from SEA-CTL to CTL-MART-1 as effector cells to screen potential candidates for their impact on target cell killing. As a positive control we used again siRNA against perforin (compare Figure 3) and analyzed transfected CTL-MART-1 by the real-time killing assay with MART-1-peptide loaded T2 cells as target cells (Figure 5A-C). Cytotoxicity was again reduced by the down-regulation of perforin (Figure 5A), however the effect was not as prominent as for SEA-CTL (compare Figure 3). The effect on the maximal killing rate was stronger compared to the effect on the endpoint target lysis (Figure 5B,C) what suggests a stronger effect in the early phase (first hour) of the killing assay. Since the down-regulation efficiency is a critical point for the screening project, we first explored the down-regulation efficiency in CTL-MART-1 for all candidates (except NMUR1 which was only expressed at very low levels). Expression of each gene was normalized to the mean of two control siRNAs transfected into CTL-MART-1. Since the down-regulation efficiency was variable for different target genes (Figure 5D), we only tested those candidates when the mRNA level of the corresponding candidate was at most 55% of the level in control conditions (24-30 hours after transfection). Consequently, we excluded TNFa, BCL2, ADRB2, CALML3, IFNG, EGR1, DMPK and EGR2 (compare Figure 5D, upper dotted line). Except these genes, the siRNAs against all other candidates were transfected into CTL-MART-1 and analyzed by the real-time killing assay. A summary of the target cell lysis, 24-48 hours after transfection, is presented in Figure 5E. Again, none of the candidates showed an average target lysis at 240 minutes higher than 115% or lower than 85%.

**Figure 5:**
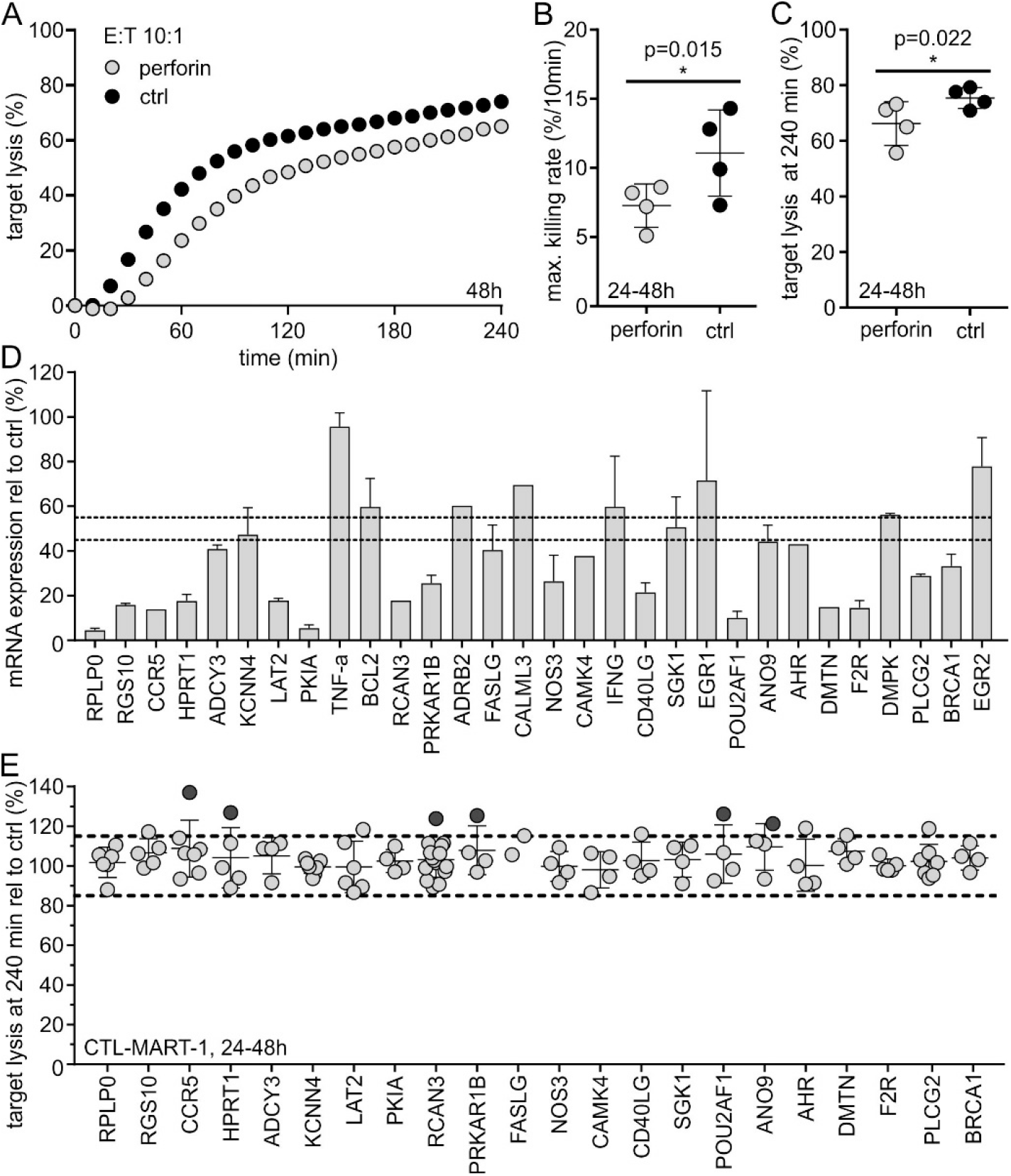
A-C. Real-time killing assay of perforin down-regulated MART-1 specific CTL (grey, CTL-MART-1) in comparison to control siRNA (mean of two control siRNAs) transfected CTL-MART-1 (black, ctrl). As target cells, MART-1 loaded T2 cells were used with an effector to target ratio (E:T) of 5:1 or 10:1. A. Representative killing traces 48h after transfection. Statistical analysis of the maximum killing rate 24h and 48h after transfection (number of transfections, n=2, B) and the endpoint target lysis 24h and 48h after transfection (number of transfections, n=2, C). Data are shown as mean +/−SD. D. Relative mRNA expression of all tested candidates 24-30 h after transfection of CTL-MART-1 with the corresponding siRNAs (CCR5, ADRB2, AHR, RCAN3, CALML3, CAMK4, DMTN, n=1; KCNN4, BCL2, n=3; all other candidates, n=2, partially shown in Figure 4B for CTL-MART1). The mRNA expression level in control siRNA transfected CTL-MART-1 was set to 100%. E. Endpoint target lysis of CTL-MART-1 transfected with siRNA against selected candidates relative to the mean of CTL-MART-1 transfected with two different control siRNAs. MART-1 loaded T2 cells were used as target cells with an E:T ratio of 5:1 or 2:1. Data are from 2-11 different transfections (n=2 for ADCY3, PKIA, PRKAR1B, FASLG, NOS3, CAMK4, CD40LG, SGK1, POU2AF1, ANO9, AHR, DMTN, DMPK and BRCA1; n=3 for RPLP0, RGS10, LAT2 and F2R; n=4 for CCR5 and PLCG2; n=6 for KCNN4; n=11 for RCAN3). Values higher than 120% target lysis at 240 min compared to the control siRNAs are labelled in dark grey. Data are shown as mean +/−SD.

Since Orai and STIM proteins are the key molecular players for Ca^2+^-entry in CTL, we reasoned that it may be difficult to identify further candidates regulating Ca^2+^-dependent cytotoxicity without putting a selective pressure on the system. Thus, in a next step, we tested several candidates under Ca^2+^-limiting conditions (by adding EGTA to the medium during the real-time killing assay, (Zhou *et al*., 2018)) based on different arguments. 1. KCNN4 and SGK1 were selected since they showed an intermediate level knockdown of target mRNAs between 45% and 55% relative to control conditions (Figure 5D). In cases of incomplete or intermediate downregulation, one can easily overlook an effect due to the low efficiency of siRNA down-regulation. 2. Candidates which showed a target cell lysis of either less than 80% or more than 120% at least in one single experiment. This criterion led to the selection of CCR5, HPRT1, RCAN3, PRKAR1B, POU2AF1 and ANO9 (Figure 5E, dark grey dots). Altogether, eight out of the originally 31 candidates were tested in a real-time killing assay under Ca^2+^-limiting conditions. To reduce the extracellular Ca^2+^ concentration ([Ca^2+^]_ext_), various concentrations of EGTA were added to the medium during the real-time killing assay. The addition of 0.80 up to 0.95 mM EGTA leads to a [Ca^2+^]_ext_ of about 10 µM to 150 µM (Zhou *et al*., 2018). Considering experiment-to-experiment variability that was observed even for CTL-MART-1 (Fig. 4A), we applied different EGTA concentrations in different experiments. To enable regulation of Ca^2+^-dependent cytotoxicity to take place in both directions (enhancement or reduction), we aimed at an intermediate cytotoxic efficiency under Ca^2+^-limiting conditions. Therefore, we always optimized the EGTA concentrations for each experiment to reach an efficiency between 30%-85% under Ca^2+^-limiting conditions relative to control conditions. Figure 6A shows examples of killing traces for control siRNA transfected CTL-MART-1 and for MART-1 peptide-loaded T2 cells as target cells with different EGTA concentrations. From these different traces, the appropriate condition was determined to be 0.93 mM EGTA because this yielded a cytotoxicity in the desired range. Choosing such a cytotoxicity range should facilitate the detection of effects of the different candidates. Based on these considerations, we tested the eight candidates according to the criteria mentioned above with the real-time killing assay under Ca^2+^-limiting conditions. As an example, Figure 6B illustrates killing traces of KCNN4 down-regulated MART-1-CTL versus control siRNA transfected cells in the defined killing range. Down-regulation of KCNN4 significantly reduced the killing capacity of MART-1-CTL by about 70% (mean target lysis at 240 min 31.2% +/−14.5% SEM) whereas down-regulation of RCAN3 enhanced the killing capacity of MART-1-CTL (mean target lysis at 240 min 119.7% +/−17.0% SEM) (Figure 6C). If the lowest “outlier” point is omitted from the analysis, the enhancement by RCAN3 down-regulation is significant (p=0.026). When we separated the assembled data for CTL-MART-1 (Figure 5E) into the two different time points, day 1 and day 2 after transfection, down-regulation of RCAN3 was significant at day 2 even without the addition of EGTA (Supplementary Figure 2). Downregulation of none of the other 6 candidates showed any significant effect, only downregulation of CCR5 led to high variations in killing efficiency at 240 minutes, a tendency which was already visible without limiting the [Ca^2+^]_ext_ (Figure 5E, Figure 6C).

**Figure 6:**
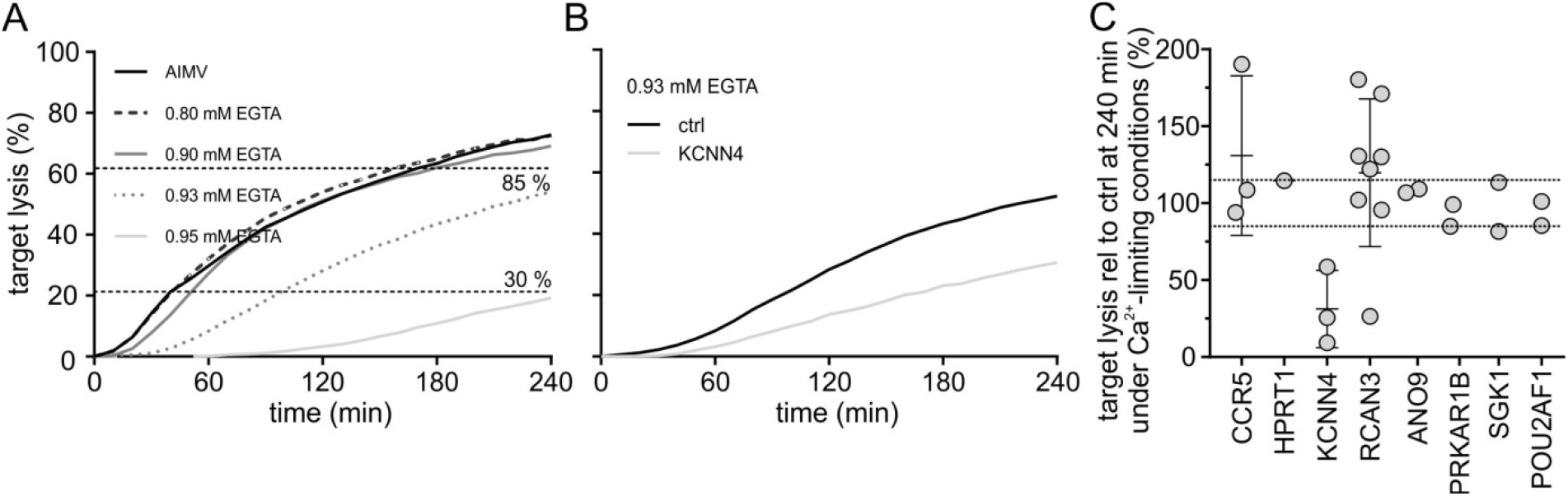
Real-time killing assay under Ca^2+^-limiting conditions manipulated by the addition of EGTA using CTL-MART-1 as effector and MART-1 peptide loaded T2 cells as target cells with an E:T ratio of 2:1. A. Representative killing traces of CTL-MART-1 transfected with a control siRNA using different concentrations of EGTA to determine the extracellular calcium concentration leading to a 30–85 % reduction in killing capacity after 240 min relative to control (defined as Ca^2+^-limiting condition). B. Representative killing traces of CTL MART-1 transfected with control siRNA (mean of two control siRNAs, black) or with KCNN4 siRNA (grey) under Ca^2+^-limiting conditions (0.93 mM EGTA) 24h after transfection. C. Endpoint target lysis of CTL-MART-1 transfected with siRNA against selected candidates relative to the mean of CTL-MART-1 transfected with two different control siRNAs. Each killing assay was performed under Ca^2+^-limiting conditions (see figure 6A) 24h to 48h after transfection. Data are from 1–6 different transfections of CTL-MART-1 (HPRT1, ANO9, PRKAR1B, SGK1 and POU2AF1, n=1; KCNN4 and CCR5, n=2; RCAN3, n=6) with 1 or 2 data points (24h and 48h after transfection). Data are shown as mean +/−SD.

Transfection with siRNA always comes at the risk of an insufficient target mRNA down-regulation (compare Figure 5D). Therefore, we also implemented an alternative approach to test for a potential function of the candidates. For 5 of our candidates, inhibitory substances are available. We tested TRAM34 on KCNN4 to confirm the down regulation in the killing efficiency seen in CTL-MART-1. Infliximab was used against TNFα, ICI118.551 against ADRB2, Maraviroc against CCR5 and Venetoclax against BCL2. The latter four ones were targeted because mRNA down-regulation of the respective mRNA was insufficient or the killing efficiency very variable (compare Figure 5D,E).

CCR5 inhibition by 10 µM Maraviroc had a slightly reducing (but significant) impact on target cell lysis (Maraviroc mean 74.2% +/−8.9% SEM and DMSO control mean 82.3% +/−8.0% SEM) (Figure 7A). 50 µM of the KCNN4 inhibitor TRAM34 reduced the target cell efficiency in CTL-MART-1 by about 40% (TRAM34 mean 47.7% +/−4.8% SEM and DMSO control mean 75.2% +/−6.7% SEM) (Figure 7B). The same tendency was also found in SEA-CTL, but this was not significant (Figure 7B). Next, we tested the inhibitors against the three candidates with insufficient mRNA down-regulation. The application of the TNFα inhibitor infliximab had no impact on the killing capacity of CTL-MART-1, neither with 1 µg/ml nor with 10 µg/ml antibody in comparison to control conditions where Rituximab, an IgG anti-CD20 antibody, was added as control (Figure 7C, left panel). When EGTA was added to the real-time killing assay to induce Ca^2+^-limiting conditions, a slight tendency towards an impairment of the killing capacity of CTL-MART-1 was detected (Figure 7C, right panel). Similarly, the application of ADRB2 inhibitor ICI 118,551 revealed only a non-significant tendency when using a high concentration of 50 µM independent of the use of Ca^2+^-limiting or non-limiting conditions (Figure 7D). The last candidate, BCL2, is inhibited by Venetoclax. In standard conditions (normal medium, AIMV, 800 µM [Ca^2+^]_ext_), 10 µM Venetoclax inhibited the target lysis at 240 min by about 33% (Venetoclax mean 60.1% +/−6.0% SEM and DMSO control mean 89.2% +/−5.5% SEM), but not at 1 µM. (Figure 7E, left panel). However application of 1 µM Venetoclax had a highly significant impact on the killing efficiency of CTL-MART-1 under Ca^2+^-limiting conditions (Venetoclax mean 40.0% +/−3.2% SEM and DMSO control mean 69.0% +/−2.3% SEM, Figure 7E) suggesting that Ca^2+^-limiting conditions unreveal the effect of BCL2 on Ca^2+^-homeostasis.

**Figure 7:**
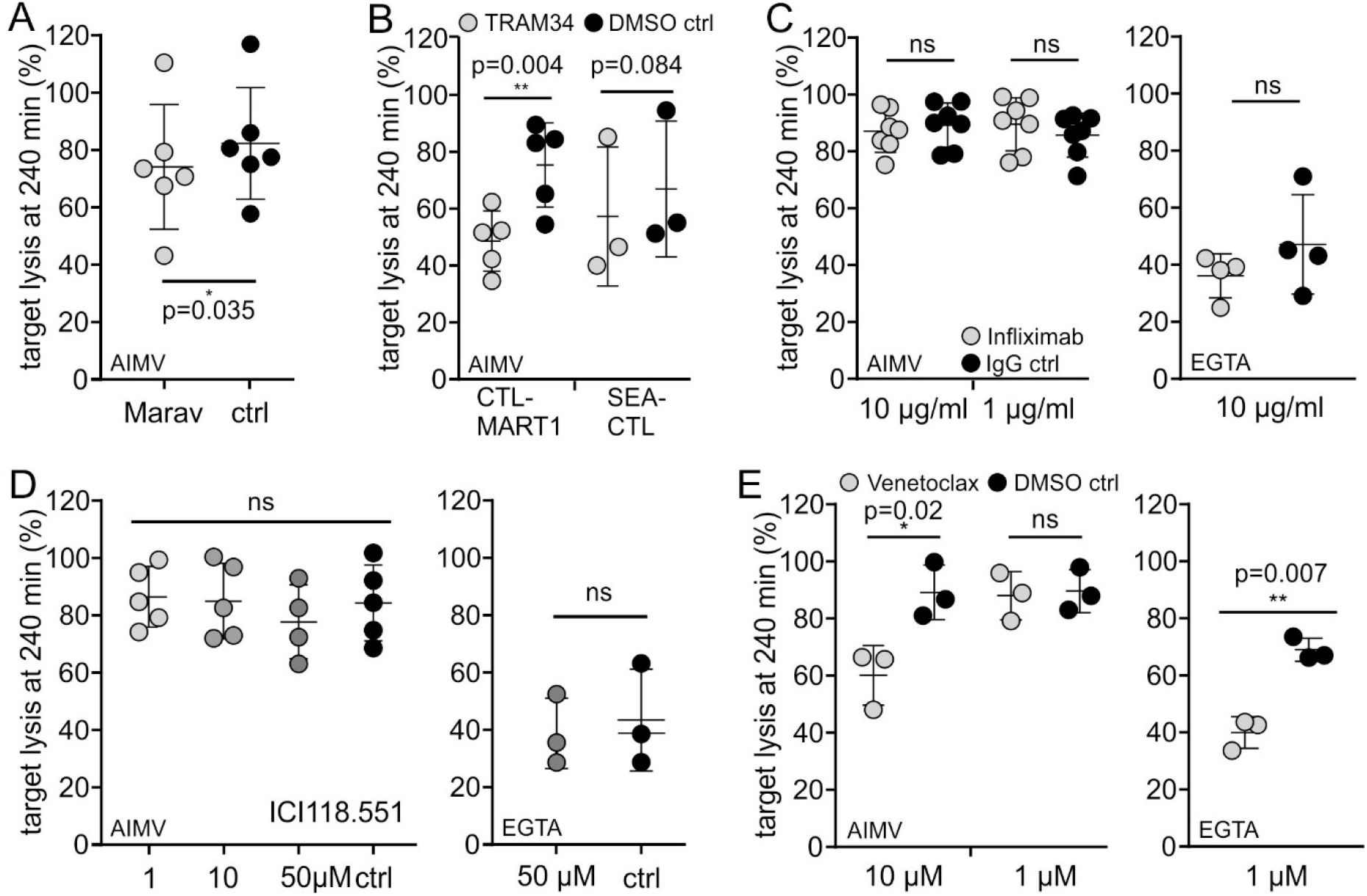
Real-time killing assay with various inhibitory substances using CTL-MART-1 as effector and MART-1 peptide loaded T2 cells as target cells with an E:T ratio of 2:1. A. Target lysis after 240 min with the application of 10 µM Maraviroc (Marav, grey circles) to inhibit CCR5 and DMSO as control (crtl, black circles). B. Target lysis after 240 min with the application of 50 µM TRAM34 (grey circles), an inhibitory substance of KCNN4, and DMSO as vehicle control (black circles). Right panel, SEA-CTL were used as effector and SEA-loaded Raji cells as target cells with an E:T ratio of 10:1. C. Target lysis after 240 min using Infliximab (grey circles), an inhibitory antibody of TNFa either in AIMV (left) or under Ca^2+^-limiting conditions (right). Rituximab was used as IgG control (black circles) in the same concentration used for Infliximab. D. Target lysis after 240 min with the application of ICI 118,551 (grey circles), inhibiting ADRB2 with different concentrations. Killing assay was performed in AIMV (left) or under Ca^2+^-limiting conditions (right). E. Target lysis after 240 min in the presence of 1 or 10 µM Venetoclax (grey circles) to inhibit the function of BCL2 in AIMV (left) and under Ca^2+^-limiting conditions manipulated by the addition of EGTA (right). Data are from 3–7 different experiments with CTL-MART-1 or from 3 different donors (SEA-CTL, B). Data are shown as mean +/−SD.

In summary, out of 31 potential candidates that we tested we identified the four proteins CCR5, KCNN4, RCAN3 and BCL2 that clearly affect the efficiency of Ca^2+^-dependent cytotoxicity of CTL-MART-1. Notably, the inhibition of CCR5, BCL2 and KCNN4 decreased cytotoxicity whereas the inhibition of RCAN3 increased the cytotoxicity.

## Discussion

Ca^2+^ dependent target cell killing of cytotoxic CD8^+^ T lymphocytes (CTL) and natural killer (NK) cells protects the human body against infection and tumor formation. Orai and STIM proteins have been identified as key components of store-operated Ca^2+^-influx. In addition, several modulators of SOCE have been published (see below). However, it is not clear how intracellular Ca^2+^ in CTL is regulated with respect to optimal killing efficiency. Thus, in this study, we screened candidates suggested from bioinformatics gene prioritization in a real-time killing assay and identified four candidates CCR5, KCNN4, RCAN3 and BCL2 modifying CTL cytotoxicity. CTL cytotoxicity is indeed of great clinical relevance in conjunction with CAR T-cell immunotherapy and identification of molecules involved in this process might have the potential to optimize the clinical outcome in the future.

### Candidates identified in this study

#### 1. KCNN4

*KCNN4* codes for the Ca^2+^-activated potassium channel K_Ca_3.1 (also known as K_Ca_3.1, IKCa^2+^ or SK4). It is a long-known and well-known ion channel in immune cells and especially in T-cells. For this reason, its function has already been extensively explored and reviewed (among others: Lewis & Cahalan, 1995; Xiao *et al*., 2003; Panyi *et al*., 2004; Feske *et al*., 2012; Trebak & Kinet, 2019). K_Ca_3.1 activity is mediated by Ca^2+^-binding to pre-bound calmodulin and the subsequent K^+^-efflux preserves the negative membrane potential as a driving force for sustained Ca^2+^-influx (Fanger *et al*., 1999). The compartmentalization of K_Ca_3.1 to the immunological synapse upon T-cell activation as a part of the signalling complex facilitates proliferation and cytokine production (Nicolaou *et al*., 2007) and also suggests a role for modulating the optimal [Ca^2+^] necessary for killing of CTL. Since the down-regulation of *KCNN4* mRNA was only about 50%, it is not surprising that we could not identify any effect on killing in SEA-CTL in culture medium (AIMV) with a free [Ca^2+^] of about 0.8 mM (Zhou *et al*., 2018). However, reducing the extracellular [Ca^2+^] down to 11 µM (corresponding to the addition of 0.93 mM EGTA, (Zhou *et al*., 2018)), we found a severe reduction in killing capacity suggesting a role for KCa3.1 in preserving optimal [Ca^2+^]. In accordance, in case extracellular [Ca^2+^] was not limited, we only found a less pronounced reduction in killing capacity using the K_Ca_3.1-inhibitor TRAM34 which blocks K_Ca_3.1 ion channel activity but not its compartmentalization within the IS (Wulff *et al*., 2000; Nicolaou *et al*., 2007). Interestingly, the effect of reducing killing capacity with the application of TRAM34 was much more pronounced in CTL-MART-1 compared to SEA-CTL (Figure 7). This is in a very good agreement with the fact that K_Ca_3.1 expression is dramatically enhanced after T-cell activation (Grissmer *et al*., 1992) and SEA-CTL are less activated compared to CTL-MART-1.

#### 2. RCAN3

In vertebrates, three regulators (RCAN, previously called DSCR, MCIP and calcipressin (CALP)) are known (Serrano-Candelas *et al*., 2014) to regulate calcineurin activity. Out of these, the interaction between RCAN3 and calcineurin is caused by one single motif (CIC motif) and inhibits the Ca^2+^-calcineurin mediated translocation of NFAT to the nucleus and thus T-cell activation (Mulero *et al*., 2007; Mulero *et al*., 2009). We observed no influence on killing in SEA-CTL in normal medium even though the down-regulation of RCAN3 mRNA was reasonably efficient (down to 30%). In CTL-MART-1, where down-regulation by siRNA was more efficient (down to 18% compared to control siRNA transfected cells), the killing efficiency in RCAN3 siRNA transfected cells was significantly enhanced in culture medium (AIMV) with a free [Ca^2+^] of about 0.8 mM (Zhou *et al*., 2018), but only 48 hours after siRNA transfection (compare Supplementary Figure 1). This suggests (which is not in a good agreement with (Klein-Hessling *et al*., 2017)) that less Ca^2+^-calcineurin mediated translocation of NFAT to the nucleus is correlated with enhanced killing even though we cannot exclude different, so far unknown modes of action, which were not the subject of this screening project.

#### 3. CCR5

The G-protein-coupled receptor, C-C chemokine receptor type 5 (CCR5) is the receptor for the β-chemokines chemokine ligand (CCL)-3 (MlP-lα), CCL4 (MIP-lβ), and CCL5 (RANTES). It is known for its central role in the entry of human immunodeficiency virus (HIV) infection into CD4^+^ T-cells, where it acts as co-receptor next to CD4 (Alkhatib *et al*., 1996; Deng *et al*., 1996). CCR5 expression in effector CD8^+^ T-cells also seems to be correlated with proliferation, with the production of inflammatory cytokines such as TNFa, interleukin 2 (IL-2) or interferon (INF)-γ (Palmer *et al*., 2010) and with the migration to inflamed tissue (Fukada *et al*., 2002). Recently, in an increased number of activated CD8^+^ T-cells, elevated expression levels of CCR5 (together with TNFR2) were identified in PBMC from glioma patients’ peripheral blood (Chen *et al*., 2019). Also CCR5 was identified as a marker for natural killer T (NKT) cells (Liu *et al*., 2019). The FDA approved CCR5 inhibitor maraviroc inhibits T-cell migration and at high concentrations (100 µM) also proliferation (Arberas *et al*., 2013). Interestingly, when inhibiting CCR5 with maraviroc in CTL-MART-1, the killing efficiency was slightly reduced which might be associated with a lower production of inflammatory cytokines.

#### 4. BCL2

Bcl-2 belongs to the Bcl-2 family proteins which are documented as regulators of apoptosis in the mitochondrial pathway existing of pro- and anti-apoptotic members (Popgeorgiev *et al*., 2018). Bcl-2 is an anti-apoptotic, pro-survival member of this family playing a central role in B-cell lymphoma development (Adams *et al*., 2018) with gene translocations in 90% of follicular lymphoma (FL) and in 20% of diffuse large B-cell lymphoma (DLBCL) (Schuetz *et al*., 2012; Kahl & Yang, 2016). In the scenario of (relapsed) chronic lymphocytic leukaemia (CLL), Bcl-2 is substantially over-expressed which leads to long-lived clonal lymphocytes (Robertson *et al*., 1996). The administration of Venetoclax, a specific inhibitor of Bcl-2, has improved outcomes for the patients (Roberts *et al*., 2016). More recently, there are also ongoing studies in which Venetoclax is administered in combination with obinutuzumab (anti-CD20 antibody) (summarized in (Salvaris & Opat, 2020)). Apart from the important role in B-cell lymphoma, Bcl-2 was also shown to directly regulate the intracellular Ca^2+^ level by inhibiting the inositol 1,4,5-trisphosphate (InsP_3_) receptor-mediated Ca^2+^-influx (Chen *et al*., 2004). Already 25 years ago, scientists discovered that Bcl-2 influences the Ca^2+^-homeostasis (also in the context of its partial localization to the mitochondrial membrane) (Baffy *et al*., 1993; Lam *et al*., 1994; Magnelli *et al*., 1994) but the direction for regulation of Ca^2+^ in the ER lumen is controversially discussed (Distelhorst & Shore, 2004). Lee and his colleagues reported a protective function of Bcl-2 in the target cells p815 against Fas- but not perforin-mediated lysis of CTL (Lee *et al*., 1996). However, to our knowledge the Bcl-2 effect on killing efficiency has not been investigated yet. Inhibition of Bcl-2 with Venetoclax significantly decreased killing efficiency of CTL-MART-1, pointing to a Ca^2+^-regulating function in these cells. Again, the molecular mode of action was not further investigated because this was not the subject of this screening project.

### Candidates which have been not identified in this study

In fact, many other molecules are known to regulate Ca^2+^-homeostasis in T-cells in general but also in CD8^+^ T-cells. Among them are: plasma membrane Ca^2+^ ATPases (PMCA), sarcoplasmic/endoplasmic reticulum Ca^2+^ ATPase (SERCA), mitochondrial Ca^2+^ uniporter (MCU), transient receptor potential (TRP) channels, purinergic ionotropic receptors (P2RXs), voltage-activated Ca^2+^ (CaV) channels, P2RXs (P2RX1, P2RX4 or P2RX7) or Piezo channels which are gated by mechanical stimuli. Many excellent and elaborate reviews summarize the function of these proteins in T-cells (Cahalan *et al*., 2001; Lewis, 2001; Feske, 2007; Oh-hora & Rao, 2008; Cahalan & Chandy, 2009; Feske *et al*., 2012; Nohara *et al*., 2015; Vaeth & Feske, 2018; Trebak & Kinet, 2019). Since we defined combined different prioritization strategies (DEG expressed between NK cells and SEA-CTL, excluding genes which were already DEG between un-stimulated CD8^+^ T-cells and SEA-CTL, reasonable expression level in SEA-CTL), we ended up with only 29 candidates. This means that we did not identify all candidates which may regulate Ca^2+^-homeostasis and therefore could also potentially regulate the killing function in NK cells or CTL.

In summary, based on modern transcriptomic analysis of microarray based expression data of CTL and NK cells, we narrowed in on a small panel of 31 candidate genes that possibly regulate the Ca^2+^-homeostasis in CTL. Due to the small number of candidates, we were able to screen them in detail in primary human cells by siRNA-directed silencing. We identified clear, reproducible effects of four candidate genes on killing efficiency when the killing assay was performed in clonal CTL-MART-1 under Ca^2+^-limiting conditions that put selective pressure on Ca^2+^-regulation. Our findings were validated in an orthogonal screen by applying small-molecule inhibitors of the candidate proteins. Hence, this study successfully identified four novel molecular players, KCNN4, RCAN3, CCR5 and BCL2, that regulate Ca^2+^ dependent killing capacity of CD8^+^ T-cells.

## Materials and Methods

### Ethical approval

The use of residual human material was approved by the local ethics committee (Ärztekammer des Saarlandes, reference 84/15, Prof. Dr. Rettig-Stürmer). Leukocyte reduction system chambers (LRSC), a byproduct of routine plateletpheresis from healthy donors, were kindly provided by the local blood bank in the Institute of Clinical Hemostaseology and Transfusion Medicine, Saarland University Medical Center. Prior to the sample being taken, blood donors provided written consent to use their blood for research purposes.

### Reagents

Calcein-AM (#C3100) was purchased from ThermoFisher Scientific, staphylococcal enterotoxin A (SEA, #S9399), Infliximab (#Y0002047), ICI 118,551 hydrochloride (#127-5MG) and Maraviroc (#PZ0002) were from Sigma Aldrich. TRAM 34 (#2946) was purchased from R&D Systems and Venetoclax (ABT-199, #ab217298) from Abcam, Rituximab (lot oN7370B16) was from the local pharmacy (Universitätsklinikum Homburg, Hoffmann La Roche). All other chemicals and reagents not specifically mentioned were from Sigma Aldrich, ThermoFisher Scientific.

### Preparation of human leukocytes and SEA-stimulated CD8^+^ T-cells

Human peripheral blood mononuclear cells (PBMC) were isolated from leukocyte reduction system chambers (LRSC) from TRIMA Accel systems after routine plateletpheresis of healthy donors at our local blood donation. Isolation of PBMC was carried out as described before (Knorck *et al*., 2018). Shortly, LRSC were rinsed with 8-10 ml Hank’s balanced salt solution (HBSS, PAA, Cölbe, Germany) followed by a standard density gradient centrifugation (leukocyte separation medium LSM 1077, PromoCell, Heidelberg, Germany). After washing, remaining erythrocytes from the PBMC layer were lysed. The pellet was again washed in HBSS and PBMC were counted (Z2 cell counter, Beckman and Coulter). Until further use PBMC were stored at 4°C. To stimulate CD8^+^ T-cells with staphylococcal enterotoxin A (SEA), 200-250 x 10^6^ PBMCs were incubated in 1,6 ml AIMV medium supplemented with 10% FCS adding 0,8 µg SEA for 1 hour (37°C, 5 % CO_2_). Afterwards cells were transferred into a 75 cm^2^ cell culture flask containing 58 ml AIMV/10% FCS and 0.05 µg/ml IL-2. After 5 days, CD8^+^ T-cells were isolated by using Dynabeads CD8 Positive isolation kit (11333D, Thermo Fisher Scientific) according to manufacturer’s instructions except for the following: PBS/0.5% BSA without EDTA as buffer 1 and 18.75 µl instead of 25 µl Dynabeads CD8 for 1 ml PBMC (10^7^ PBMC) were used. Isolated CD8^+^ T-cells were cultured in complete AIMV medium with 0.05 µg/ml SEA added until further use.

### Cells

Raji (ATCC CCL-86), T2 target cells and Epstein-Barr Virus-transformed B lymphoblastoid cell lines (EBV-LCLs) cells were cultured in RPMI-1640 (Thermo Fisher Scientific) supplemented with 10% FCS and 1% penicillin/streptomycin (#P433, Thermo Fisher Scientific) at 37°C and 5% CO_2_. MART-1-specific CD8^+^ T-cell clones (CTL-MART-1) were generated and expanded as described previously (Friedmann *et al*., 2020). The MART-1-specific clone CTL-M3 from different expansions was used through the study (Friedmann *et al*., 2020).

### Transfection of siRNA

Transfection of siRNA was carried out using the 4D Nucleofector (Lonza) following the manufacturer’s instructions. For each transfection, 5-6 x 10^6^ SEA-stimulated CTL (SEA-CTL) or MART-1 antigen-specific T-cell clones (CTL-MART-1) were washed in PBS/0.5%BSA and resuspended in 100 µl Nucleofector solution (Amaxa™ P3 Primary Cell 4D Nucleofector TM X Kit L, Lonza) with 4-6 µl of the appropriate siRNA (20-40 µM) added. Following nucleofection (program EO115), cells were cultured in AIMV/10% FCS for at least 6 hours before adding 0.01 µg/ml IL-2. SiRNAs were purchased from Dharmacon (a part of Horizon) and are listed in table 1.

### Real-time killing assay

Cytotoxicity of siRNA-transfected SEA-stimulated CTL or antigen-specific T-cell clones was measured with a real-time killing assay over a period of 4 hours every 10 minutes (Kummerow *et al*., 2014) using Raji or antigen (MART-1) pulsed T2 cells as target cells, respectively. Raji cells were pulsed with 1 µg/ml SEA in AIMV medium + 10% FCS for 30 min and T2 cells with 5 µg/ml MART-1 peptide. Subsequently, target cells were loaded with calcein-AM (500 nM, #C3100MP, Thermo Fisher Scientific) for 15 min, washed with AIMV medium without FCS and settled at a number of 2.5×10^4^ cells per well into a 96-well black plate with clear bottom (#734-2480, VWR) for around 30 minutes. Next, transfected SEA-stimulated CTL or MART-1-T-cell clones were added to the corresponding target cells at the indicated effector to target ratio. Target lysis was measured either in the Genios Pro (Tecan, Crailsheim, Germany) or in the Clariostar (BMG Labtech, Ortenberg, Germany) using the bottom reading function at 37°C. The real-time killing assay under Ca^2+^-reduced conditions was performed as described in (Zhou *et al*., 2018). To study the impact of Maraviroc (20 mM stock solution in DMSO), Venetoclax (10 mM stock solution in DMSO), TRAM34 (10 mM stock solution in DMSO), Infliximab (35,6 mg/ml stock solution in dH_2_O) or ICI 118,551 (10 mM stock solution in dH_2_O) on cytotoxicity, SEA-CTL or CTL-MART-1 were pre-incubated for 90 min in AIMV and 10 mM Hepes containing the indicated concentration of the corresponding inhibitor or as control the corresponding concentration of DMSO, when solvated in DMSO or in case of infliximab the corresponding concentration of IgG (Rituximab, 10 mg/ml stock solution). The killing assay was performed as described above. The indicated concentration of each inhibitor was present during the complete experiment.

### RNA isolation and qRT-PCR

Experiments were done as described in (Wenning *et al*., 2011). Briefly, total RNA was isolated from either 1.5×10^6^ SEA stimulated CTL or MART-1-specific CD8^+^ T-cell clones using TRIzol reagent (Thermo Fisher Scientific) containing 1 µl Glycogen (5 µg/µl, Thermo Fisher Scientific). 0.8 µg total RNA was used for reverse transcription and 0.5 µl of cDNA was used for quantitative real time polymerase chain reaction (qRT-PCR). qRT-PCR was carried out in a CFX96™ Real-Time System C1000™ Thermal Cycler (Software Biorad CFX Manager, Version 3.0) using QuantiTect SYBR Green PCR Kit (Qiagen, #204145). QuantiTect Primer Assays were purchased from Qiagen. Primer information is summarized in table 2.

**Table 2:** Primers used in this study. Primers for the reference genes RNAPol (NM_000937) and TBP (NM_003194) are taken from (Wenning *et al*., 2011) and primers for perforin (NM_005041 and NM_001083116) from (Bhat *et al*., 2016). Primers for FASLG (NM_000639) forward 5’ GCACACAGCATCATCTTTGG 3’ and reverse 5’ CAAGATTGACCCCGGAAGTA 3’.

| gene |  | QuantiTect<br>Primer Assay |
| --- | --- | --- |
| ADCY3 | Hs_ADCY3_1_SG | QT00075348 |
| ADRB2 | Hs_ADRB2_1_SG | QT00200011 |
| AHR | Hs_AHR_2_SG | QT02422938 |
| ANO9 | Hs_ANO9_1_SG | QT01322643 |
| BCL2 | Hs_BCL2_1_SG | QT00025011 |
| BRCA1 | Hs_BRCA1_1_SG | QT00039305 |
| CALML3 | Hs_CALML3_1_SG | QT00214935 |
| CAMK4 | Hs_CAMK4_1_SG | QT00038101 |
| CCR5 | Hs_CCR5_1_SG | QT00998802 |
| CD40LG | Hs_CD40LG_1_SG | QT00000343 |
| DMPK | Hs_DMPK_1_SG | QT00094199 |
| DMTN | Hs_DMTN_1_SG | QT00493052 |
| Egr1 | Hs_EGR1_1_SG | QT00218505 |
| Egr2 | Hs_EGR2_1_SG | QT00000924 |
| F2R | Hs_F2R_1_SG | QT00230489 |
| HPRT1 | Hs_HPRT1_1_SG | QT00059066 |
| IFNG | Hs_IFNG_1_SG | QT00000525 |
| KCNN4 | Hs_KCNN4_1_SG | QT00003780 |
| LAT2 | Hs_LAT2_1_SG | QT00010934 |
| NMUR1 | Hs_NMUR1_1_SG | QT00080311 |
| NOS3 | Hs_NOS3_1_SG | QT00089033 |
| PKIA | Hs_PKIA_2_SG | QT02289658 |
| PLCG2 | Hs_PLCG2_1_SG | QT00050393 |
| POU2AF1 | Hs_POU2AF1_1_SG | QT00001540 |
| PRKAR1B | Hs_PRKAR1B_2_SG | QT02306521 |
| RCAN3 | Hs_RCAN3_1_SG | QT00075369 |
| RGS10 | Hs_RGS10_1_SG | QT00235900 |
| RPLP0 | Hs_RPLP0_1_SG | QT00075012 |
| SGK1 | Hs_SGK1_1_SG | QT00041293 |
| TNFA | Hs_TNF_1_SG | QT00029162 |

### Data and statistical analysis

Data were tested for significance using Student t-test whenever Gaussian distribution was confirmed. Otherwise, the non-parametric Wilcoxon-Mann-Whitney test was used. * p<0.05; ** p<0.01; *** p<0.001; ns, no significant difference; as stated in the figure legends. Data analyses were performed using Microsoft Excel 2016, and GraphPad Prism 7 software.

### Microarray experiments

RNA processing, purity and integrity of the input template RNA and microarray experiments were done as described before (Bhat *et al*., 2016). Microarray data are available at the GEO Platform (https://www.ncbi.nlm.nih.gov/geo/) under the accession number GSE168692.

### Data pre-processing

The raw data were normalized using quantile normalization and log2 transformed with the statistical programming language R (Ihaka & Gentleman, 1996). Expression profiles of probe sets belonging to single genes were summarized/aggregated by computing the mean expression values as reported previously in (Hamed *et al*., 2015). The normalized summarized/aggregated data were further subjected to differential expression analysis using the moderated t-test to identify the differentially expressed genes (DEGs) between the various sample groups. Namely, genes that exhibited 2-fold changes and p-value below a cutoff of 0.05 were classified as differentially expressed genes (DEGs). P-values were adjusted using the Benjamini-Hochberg (Hochberg & Benjamini, 1990) procedure to limit the false discovery rate to 5%.

### Identification of candidate genes

Candidate genes that are possibly responsible for modulating the killing efficiency of SEA-CTL are assumed to show differential expression between SEA-CTL (CD8^+^SEA) and NK samples. These are termed SEA-DEGs (differentially expressed genes). From this list, we deleted genes that are also differentially expressed between “SEA”-stimulated cells and “UN”-stimulated cells. This yielded 2349 SEA-DEGs between CD8^+^SEA and NK samples. We then used two strategies for pruning this set of differentially expressed genes to a manageable number of candidate genes.

A list of 512 Calcium-associated genes was compiled from the KEGG pathway (Kanehisa *et al*., 2016), the Gene Ontology (The Gene Ontology, 2019), the Wikipathways (Martens *et al*., 2020), and the PathwaysFinder (Yao *et al*., 2004) databases.

**Strategy 1**: The list of 2349 SEA-DEGs was intersected with the 512 “Ca^2+^-associated genes” and this resulted in 61 Ca^2+^-associated DEGs. From this list we removed genes having low expression levels in SEA-CTL (2^7.2^ (fluorescence value < 147), the mean expression value). After this, 29 candidate genes remained.

**Strategy 2**: This strategy followed that of our method TopControl (Nazarieh & Helms, 2019). First, the set of 2349 SEA-DEGs was provided as input to the webserver TFmiR (Hamed *et al*., 2015). TFmiR constructed a differential co-regulatory network involving 98 genes (transcription factors or target genes) and identified the set of 10% most connected hub genes. In this network, we identified a minimum dominating set (MDS) of genes using the instruction level parallelism (ILP) approach. In the largest connected component underlying the undirected graph of this network, we identified a minimum connected dominating set (MCDS) using the heuristic approach. Both algorithms are described in (Nazarieh *et al*., 2016). 47 genes belonged to at least one of these three sets (hubs, MDS, MCDS). Intersecting this set of 47 genes with the 512 “Calcium-associated genes” gave 7 candidate genes. Of these 7 candidates, the gene ATF3 was excluded due to its low expression level in SEA-CTL, and four other genes: Bcl2, TNF, FASLG, and EGR1 were identified by both strategies.

The union of both strategies gave 31 unique candidate genes that were further investigated.

## Additional information

### Competing interests

The authors declare no competing or conflicting interests.

### Author contributions

S.Z., G.S., V.K. and C.H. designed and performed experiments. Mo.H. and M.N. performed the data pre-processing, the computational framework and analyzed the data. S.Z., Ma.H., Mo.H., V.H. and E.C.S. designed the study. E.M. contributed to the microarray experiments. S.Z. and E.C.S. coordinated data analyses and designed final figure layout. M.N. contributed Supplementary Figure 1. E.C.S. wrote the manuscript in consultation with S.Z., Ma.H. and V.H and contribution of Mo.H. and M.N.. Authors carefully checked the manuscript and provided critical feedback.

### Funding

This work was funded by the Deutsche Forschungsgemeinschaft (DFG, the collaborative research centers SFB 1027 (project A2 to MH, project C3 to VH) and SFB 894 (project A1 to MH)).

### Data Availability Statement

Microarray data supporting the findings of this study are openly available at the GEO Platform (https://www.ncbi.nlm.nih.gov/geo/) under the accession number GSE168692.

## Acknowledgements

We are very grateful to Petra Leidinger-Kaufmann (Human Genetics, Saarland University, Homburg) for performing the microarray measurement. We thank PD Dr. Frank Neumann (José Carreras Center, Saarland University Medical Center, Homburg, Germany) for providing T2 target cells and Epstein-Barr Virus-transformed B lymphoblastoid cell lines (EBV-LCLs). We thank Carmen Hässig for cell preparation. We also thank Sandra Janku for language proof reading. We thank Prof. Dr. Hermann Eichler, Dr. Jürgen Groß and his co-workers (Institute of Clinical Hemostaseology and Transfusion Medicine, Saarland University Medical Center) for providing LRS chambers and all human blood donors.

## Notes

https://www.ncbi.nlm.nih.gov/geo/query/acc.cgi?acc=GSE168692

### Competing Interest Statement

The authors have declared no competing interest.

### Summary of Updates

Section Manuscript Basics The Article Category was changed to "New results" as it was in the first submission. With the remission it was changed to "confirmatory results" by mistake.

## References

Adams CM, Clark-Garvey S, Porcu P & Eischen CM. (2018). Targeting the Bcl-2 Family in B Cell Lymphoma. Front Oncol 8, 636.

Alkhatib G, Combadiere C, Broder CC, Feng Y, Kennedy PE, Murphy PM & Berger EA. (1996). CC CKR5: a RANTES, MIP-1alpha, MIP-1beta receptor as a fusion cofactor for macrophage-tropic HIV-1. Science 272, 1955–1958.

Arberas H, Guardo AC, Bargallo ME, Maleno MJ, Calvo M, Blanco JL, Garcia F, Gatell JM & Plana M. (2013). In vitro effects of the CCR5 inhibitor maraviroc on human T cell function. J Antimicrob Chemother 68, 577–586.

Badou A, Jha MK, Matza D & Flavell RA. (2013). Emerging roles of L-type voltage-gated and other calcium channels in T lymphocytes. Front Immunol 4, 243.

Baffy G, Miyashita T, Williamson JR & Reed JC. (1993). Apoptosis induced by withdrawal of interleukin-3 (IL-3) from an IL-3-dependent hematopoietic cell line is associated with repartitioning of intracellular calcium and is blocked by enforced Bcl-2 oncoprotein production. J Biol Chem 268, 6511–6519.

Bellucci R, Nguyen HN, Martin A, Heinrichs S, Schinzel AC, Hahn WC & Ritz J. (2012). Tyrosine kinase pathways modulate tumor susceptibility to natural killer cells. J Clin Invest 122, 2369–2383.

Bhat SS, Friedmann KS, Knorck A, Hoxha C, Leidinger P, Backes C, Meese E, Keller A, Rettig J, Hoth M, Qu B & Schwarz EC. (2016). Syntaxin 8 is required for efficient lytic granule trafficking in cytotoxic T lymphocytes. Biochimica et biophysica acta 1863, 1653–1664.

Cahalan MD & Chandy KG. (2009). The functional network of ion channels in T lymphocytes. Immunol Rev 231, 59–87.

Cahalan MD, Wulff H & Chandy KG. (2001). Molecular properties and physiological roles of ion channels in the immune system. J Clin Immunol 21, 235–252.

Chen PY, Wu CY, Fang JH, Chen HC, Feng LY, Huang CY, Wei KC, Fang JY & Lin CY. (2019). Functional Change of Effector Tumor-Infiltrating CCR5(+)CD38(+)HLA-DR(+)CD8(+) T Cells in Glioma Microenvironment. Front Immunol 10, 2395.

Chen R, Valencia I, Zhong F, McColl KS, Roderick HL, Bootman MD, Berridge MJ, Conway SJ, Holmes AB, Mignery GA, Velez P & Distelhorst CW. (2004). Bcl-2 functionally interacts with inositol 1,4,5-trisphosphate receptors to regulate calcium release from the ER in response to inositol 1,4,5-trisphosphate. J Cell Biol 166, 193–203.

Deng H, Liu R, Ellmeier W, Choe S, Unutmaz D, Burkhart M, Di Marzio P, Marmon S, Sutton RE, Hill CM, Davis CB, Peiper SC, Schall TJ, Littman DR & Landau NR. (1996). Identification of a major co-receptor for primary isolates of HIV-1. Nature 381, 661–666.

Distelhorst CW & Shore GC. (2004). Bcl-2 and calcium: controversy beneath the surface. Oncogene 23, 2875–2880.

Fanger CM, Ghanshani S, Logsdon NJ, Rauer H, Kalman K, Zhou J, Beckingham K, Chandy KG, Cahalan MD & Aiyar J. (1999). Calmodulin mediates calcium-dependent activation of the intermediate conductance KCa channel, IKCa1. J Biol Chem 274, 5746–5754.

Fenninger F & Jefferies WA. (2019). What’s Bred in the Bone: Calcium Channels in Lymphocytes. J Immunol 202, 1021–1030.

Feske S. (2007). Calcium signalling in lymphocyte activation and disease. Nat Rev Immunol 7, 690–702.

Feske S. (2013). Ca(2+) influx in T cells: how many ca(2+) channels? Front Immunol 4, 99.

Feske S, Skolnik EY & Prakriya M. (2012). Ion channels and transporters in lymphocyte function and immunity. Nat Rev Immunol 12, 532–547.

Friedmann KS, Bozem M & Hoth M. (2019). Calcium signal dynamics in T lymphocytes: Comparing in vivo and in vitro measurements. Semin Cell Dev Biol 94, 84–93.

Friedmann KS, Knörck A, Cappello S, Hoxha C, Schwär G, Iden S, Bogeski I, Kummerow C, Schwarz EC & Hoth M. (2020). Combined CTL and NK cell cytotoxicity against cancer cells. bioRxiv, 2020.2006.2014.150672.

Fukada K, Sobao Y, Tomiyama H, Oka S & Takiguchi M. (2002). Functional expression of the chemokine receptor CCR5 on virus epitope-specific memory and effector CD8+ T cells. J Immunol 168, 2225–2232.

Grissmer S, Lewis RS & Cahalan MD. (1992). Ca(2+)-activated K+ channels in human leukemic T cells. J Gen Physiol 99, 63–84.

Hamed M, Spaniol C, Zapp A & Helms V. (2015). Integrative network-based approach identifies key genetic elements in breast invasive carcinoma. BMC Genomics 16 Suppl 5, S2.

Hattab D & Bakhtiar A. (2020). Bioengineered siRNA-Based Nanoplatforms Targeting Molecular Signaling Pathways for the Treatment of Triple Negative Breast Cancer: Preclinical and Clinical Advancements. Pharmaceutics 12.

Hochberg Y & Benjamini Y. (1990). More powerful procedures for multiple significance testing. Stat Med 9, 811–818.

Ihaka R & Gentleman R. (1996). R: A Language for Data Analysis and Graphics. Journal of Computational and Graphical Statistics 5, 299–314.

Kahl BS & Yang DT. (2016). Follicular lymphoma: evolving therapeutic strategies. Blood 127, 2055–2063.

Kanehisa M, Sato Y, Kawashima M, Furumichi M & Tanabe M. (2016). KEGG as a reference resource for gene and protein annotation. Nucleic Acids Res 44, D457–462.

Khandelwal N, Breinig M, Speck T, Michels T, Kreutzer C, Sorrentino A, Sharma AK, Umansky L, Conrad H, Poschke I, Offringa R, Konig R, Bernhard H, Machlenkin A, Boutros M & Beckhove P. (2015). A high-throughput RNAi screen for detection of immune-checkpoint molecules that mediate tumor resistance to cytotoxic T lymphocytes. EMBO Mol Med 7, 450–463.

Klein-Hessling S, Muhammad K, Klein M, Pusch T, Rudolf R, Floter J, Qureischi M, Beilhack A, Vaeth M, Kummerow C, Backes C, Schoppmeyer R, Hahn U, Hoth M, Bopp T, Berberich-Siebelt F, Patra A, Avots A, Muller N, Schulze A & Serfling E. (2017). NFATc1 controls the cytotoxicity of CD8(+) T cells. Nat Commun 8, 511.

Knorck A, Marx S, Friedmann KS, Zophel S, Lieblang L, Hassig C, Muller I, Pilch J, Sester U, Hoth M, Eichler H, Sester M & Schwarz EC. (2018). Quantity, quality, and functionality of peripheral blood cells derived from residual blood of different apheresis kits. Transfusion 58, 1516–1526.

Kummerow C, Schwarz EC, Bufe B, Zufall F, Hoth M & Qu B. (2014). A simple, economic, time-resolved killing assay. Eur J Immunol 44, 1870–1872.

Lam M, Dubyak G, Chen L, Nunez G, Miesfeld RL & Distelhorst CW. (1994). Evidence that BCL-2 represses apoptosis by regulating endoplasmic reticulum-associated Ca2+ fluxes. Proc Natl Acad Sci U S A 91, 6569–6573.

Lee RK, Spielman J & Podack ER. (1996). Bcl-2 protects against Fas-based but not perforin-based T cell-mediated cytolysis. Int Immunol 8, 991–1000.

Lewis RS. (2001). Calcium signaling mechanisms in T lymphocytes. Annu Rev Immunol 19, 497–521.

Lewis RS & Cahalan MD. (1995). Potassium and calcium channels in lymphocytes. Annu Rev Immunol 13, 623–653.

Liu J, Hill BJ, Darko S, Song K, Quigley MF, Asher TE, Morita Y, Greenaway HY, Venturi V, Douek DC, Davenport MP, Price DA & Roederer M. (2019). The peripheral differentiation of human natural killer T cells. Immunol Cell Biol 97, 586–596.

Magnelli L, Cinelli M, Turchetti A & Chiarugi VP. (1994). Bcl-2 overexpression abolishes early calcium waving preceding apoptosis in NIH-3T3 murine fibroblasts. Biochem Biophys Res Commun 204, 84–90.

Mainini F & Eccles MR. (2020). Lipid and Polymer-Based Nanoparticle siRNA Delivery Systems for Cancer Therapy. Molecules 25.

Mantei A, Rutz S, Janke M, Kirchhoff D, Jung U, Patzel V, Vogel U, Rudel T, Andreou I, Weber M & Scheffold A. (2008). siRNA stabilization prolongs gene knockdown in primary T lymphocytes. Eur J Immunol 38, 2616–2625.

Martens M, Ammar A, Riutta A, Waagmeester A, Slenter DN, Hanspers K, R AM, Digles D, Lopes EN, Ehrhart F, Dupuis LJ, Winckers LA, Coort SL, Willighagen EL, Evelo CT, Pico AR & Kutmon M. (2020). WikiPathways: connecting communities. Nucleic Acids Res.

Maul-Pavicic A, Chiang SC, Rensing-Ehl A, Jessen B, Fauriat C, Wood SM, Sjoqvist S, Hufnagel M, Schulze I, Bass T, Schamel WW, Fuchs S, Pircher H, McCarl CA, Mikoshiba K, Schwarz K, Feske S, Bryceson YT & Ehl S. (2011). ORAI1-mediated calcium influx is required for human cytotoxic lymphocyte degranulation and target cell lysis. Proc Natl Acad Sci U S A 108, 3324–3329.

Moffat J & Sabatini DM. (2006). Building mammalian signalling pathways with RNAi screens. Nat Rev Mol Cell Biol 7, 177–187.

Mulero MC, Aubareda A, Orzaez M, Messeguer J, Serrano-Candelas E, Martinez-Hoyer S, Messeguer A, Perez-Paya E & Perez-Riba M. (2009). Inhibiting the calcineurin-NFAT (nuclear factor of activated T cells) signaling pathway with a regulator of calcineurin-derived peptide without affecting general calcineurin phosphatase activity. J Biol Chem 284, 9394–9401.

Mulero MC, Aubareda A, Schluter A & Perez-Riba M. (2007). RCAN3, a novel calcineurin inhibitor that down-regulates NFAT-dependent cytokine gene expression. Biochimica et biophysica acta 1773, 330–341.

Nazarieh M & Helms V. (2019). TopControl: A Tool to Prioritize Candidate Disease-associated Genes based on Topological Network Features. Scientific reports 9, 19472.

Nazarieh M, Wiese A, Will T, Hamed M & Helms V. (2016). Identification of key player genes in gene regulatory networks. BMC Syst Biol 10, 88.

Nicolaou SA, Neumeier L, Peng Y, Devor DC & Conforti L. (2007). The Ca(2+)-activated K(+) channel KCa3.1 compartmentalizes in the immunological synapse of human T lymphocytes. Am J Physiol Cell Physiol 292, C1431–1439.

Nohara LL, Stanwood SR, Omilusik KD & Jefferies WA. (2015). Tweeters, Woofers and Horns: The Complex Orchestration of Calcium Currents in T Lymphocytes. Front Immunol 6, 234.

Oh-hora M & Rao A. (2008). Calcium signaling in lymphocytes. Curr Opin Immunol 20, 250–258.

Palmer LA, Sale GE, Balogun JI, Li D, Jones D, Molldrem JJ, Storb RF & Ma Q. (2010). Chemokine receptor CCR5 mediates alloimmune responses in graft-versus-host disease. Biol Blood Marrow Transplant 16, 311–319.

Panyi G, Varga Z & Gaspar R. (2004). Ion channels and lymphocyte activation. Immunol Lett 92, 55–66.

Pelletier L & Savignac M. (2013). Ca(2+) signaling in T-cell subsets with a focus on the role of cav1 channels: possible implications in therapeutics. Front Immunol 4, 150.

Popgeorgiev N, Jabbour L & Gillet G. (2018). Subcellular Localization and Dynamics of the Bcl-2 Family of Proteins. Front Cell Dev Biol 6, 13.

Roberts AW, Davids MS, Pagel JM, Kahl BS, Puvvada SD, Gerecitano JF, Kipps TJ, Anderson MA, Brown JR, Gressick L, Wong S, Dunbar M, Zhu M, Desai MB, Cerri E, Heitner Enschede S, Humerickhouse RA, Wierda WG & Seymour JF. (2016). Targeting BCL2 with Venetoclax in Relapsed Chronic Lymphocytic Leukemia. N Engl J Med 374, 311–322.

Robertson LE, Plunkett W, McConnell K, Keating MJ & McDonnell TJ. (1996). Bcl-2 expression in chronic lymphocytic leukemia and its correlation with the induction of apoptosis and clinical outcome. Leukemia 10, 456–459.

Salvaris R & Opat S. (2020). An update of venetoclax and obinutuzumab in chronic lymphocytic leukemia. Future Oncol.

Schuetz JM, Johnson NA, Morin RD, Scott DW, Tan K, Ben-Nierah S, Boyle M, Slack GW, Marra MA, Connors JM, Brooks-Wilson AR & Gascoyne RD. (2012). BCL2 mutations in diffuse large B-cell lymphoma. Leukemia 26, 1383–1390.

Serrano-Candelas E, Farre D, Aranguren-Ibanez A, Martinez-Hoyer S & Perez-Riba M. (2014). The vertebrate RCAN gene family: novel insights into evolution, structure and regulation. PLoS One 9, e85539.

Shih T, De S & Barnes BJ. (2019). RNAi Transfection Optimized in Primary Naive B Cells for the Targeted Analysis of Human Plasma Cell Differentiation. Front Immunol 10, 1652.

Sioud M. (2020). Optimized siRNA Delivery into Primary Immune Cells Using Electroporation. Methods Mol Biol 2115, 119–131.

Smith N, Vidalain PO, Nisole S & Herbeuval JP. (2016). An efficient method for gene silencing in human primary plasmacytoid dendritic cells: silencing of the TLR7/IRF-7 pathway as a proof of concept. Scientific reports 6, 29891.

The Gene Ontology C. (2019). The Gene Ontology Resource: 20 years and still GOing strong. Nucleic Acids Res 47, D330–D338.

Trebak M & Kinet JP. (2019). Calcium signalling in T cells. Nat Rev Immunol 19, 154–169.

Vaeth M & Feske S. (2018). Ion channelopathies of the immune system. Curr Opin Immunol 52, 39–50.

Wenning AS, Neblung K, Strauss B, Wolfs MJ, Sappok A, Hoth M & Schwarz EC. (2011). TRP expression pattern and the functional importance of TRPC3 in primary human T-cells. Biochimica et biophysica acta 1813, 412–423.

Wulff H, Miller MJ, Hansel W, Grissmer S, Cahalan MD & Chandy KG. (2000). Design of a potent and selective inhibitor of the intermediate-conductance Ca2+-activated K+ channel, IKCa1: a potential immunosuppressant. Proc Natl Acad Sci U S A 97, 8151–8156.

Xiao L, Fu HY, Song DM & Fan SG. (2003). [Ion channels on T lymphocyte]. Sheng Li Ke Xue Jin Zhan 34, 105–110.

Yao D, Wang J, Lu Y, Noble N, Sun H, Zhu X, Lin N, Payan DG, L. M & Qu K. (2004). PathwayFinder: paving the way towards automatic pathway extraction. In Proceedings of the second conference on Asia-Pacific bioinformatics - Volume 29, pp. 53–62. Australian Computer Society, Inc., Dunedin, New Zealand.

Zhou LY, Qin Z, Zhu YH, He ZY & Xu T. (2019). Current RNA-based Therapeutics in Clinical Trials. Curr Gene Ther 19, 172–196.

Zhou P, Shaffer DR, Alvarez Arias DA, Nakazaki Y, Pos W, Torres AJ, Cremasco V, Dougan SK, Cowley GS, Elpek K, Brogdon J, Lamb J, Turley SJ, Ploegh HL, Root DE, Love JC, Dranoff G, Hacohen N, Cantor H & Wucherpfennig KW. (2014). In vivo discovery of immunotherapy targets in the tumour microenvironment. Nature 506, 52–57.

Zhou X, Friedmann KS, Lyrmann H, Zhou Y, Schoppmeyer R, Knorck A, Mang S, Hoxha C, Angenendt A, Backes CS, Mangerich C, Zhao R, Cappello S, Schwar G, Hassig C, Neef M, Bufe B, Zufall F, Kruse K, Niemeyer BA, Lis A, Qu B, Kummerow C, Schwarz EC & Hoth M. (2018). A calcium optimum for cytotoxic T lymphocyte and natural killer cell cytotoxicity. The Journal of physiology 596, 2681–2698.

