## Supplementary Figure 1 and Figure 2 for "Identification of molecular candidates which regulate calcium-dependent CD8^+^ T-cell cytotoxicity"

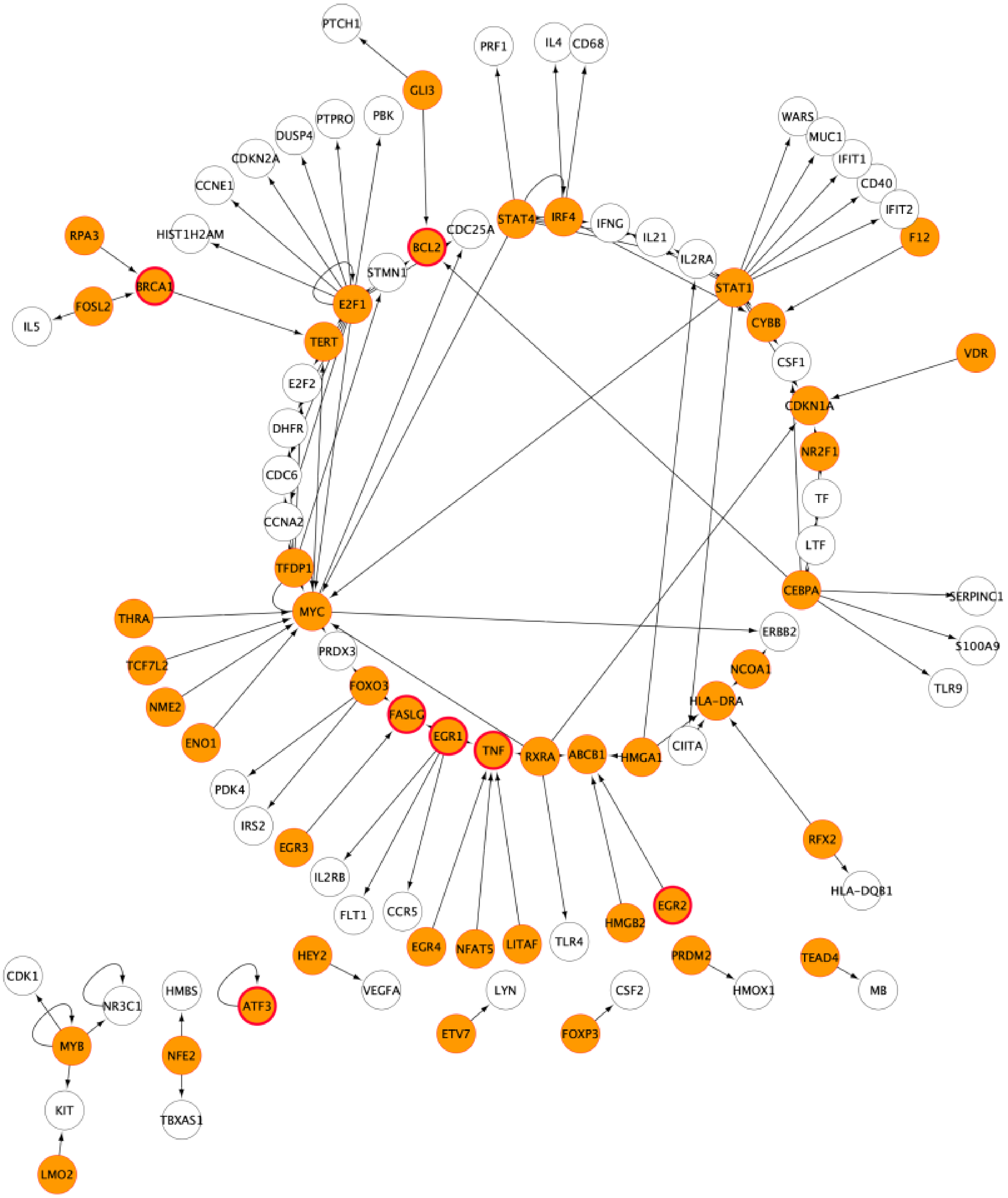


**Supplementary Figure 1**: Regulatory interactions (pointed arcs) detected by the TFmiR webserver involving the deregulated genes. The network consists of 98 nodes distributed in 9 components with 79 in the largest connected component displayed in a circular layout. Marked in orange are genes that play a role in the network either as a high-degree central node and/or as a member in the MDS or MCDS sets. Marked in orange on the inner circle are genes that have a high degree of centrality (E2F1, MYC, STAT1, CEBPA, IRF4, STAT4, TFDP1, EGR1, RXRA) in addition to being part of MDS and/or MCDS. Genes labeled by a red circle border are part of $Ca^{2+}$-associated pathways.

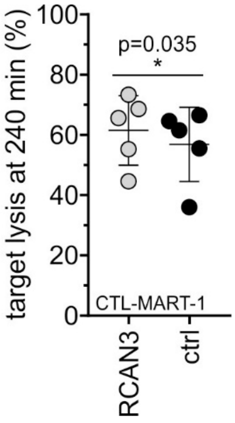


**Supplementary Figure 2**: Real-time killing assay using CTL-MART-1 as effector and MART-1 peptide loaded T2 cells as target cells with an E:T ratio of 2:1. Target lysis after 240 min is shown in RCAN3 or the mean of two control siRNA transfected CTL-MART-1 48h after transfection. Data are from 4 different transfections of CTL-MART-1. Data are shown as mean +/- SD.
